# Nanoscale Molecular Characterisation of Hair Cuticles using Integrated AFM-IR

**DOI:** 10.1101/2020.02.12.946103

**Authors:** A. P. Fellows, M. T. L. Casford, P. B. Davies

**Affiliations:** University of Cambridge

**Keywords:** Cuticle sub-structure, AFM-IR, nanometre IR mapping, protein, lipids, hair

## Abstract

The nanometre-scale topography and chemical structure of hair cuticles has been investigated by vibrational spectroscopy and imaging in two spectral regions. The combination of Atomic Force Microscopy with a tuneable infrared laser (AFM-IR) circumvents the diffraction limit that has impaired traditional infrared spectroscopy, facilitating surface spectroscopy at ultra-spatial resolution. The variation in protein and lipid content of the cuticle cell surface approaching its edge, as well as the exposed layered structure of the cell at the edge itself, was investigated. Furthermore, the contribution of cystine-related products to the cuticle layers was determined. The variation of protein, lipid and cystine composition in the observed layers, as well as the measured dimensions of each, correspond closely to that of the epicuticle, A-layer, exocuticle and endocuticle layers of the cuticle cell sub-structure.

**Statement of Significance:** Using AFM-IR to analyse the nanoscale cuticle features is both significant and novel in the field. Thus far, the great majority of work on the chemical investigation of the structure of hair has been limited to bulk measurements, or subject to the diffraction limit associated with traditional IR spectroscopies and microscopies. AFM-IR circumvents this diffraction limit and allows nanometre-scale, localised chemical investigation with high surface selectivity. While non-chemical investigations, e.g. those using Transmission Election Microscopy, have previously shown cuticles to have a layered substructure, AFM-IR sheds light on significant chemical variations of protein and lipid compositions within such layers, enabling their quantification.

## Introduction

The primary importance of hair for mammals is in providing thermal insulation. Additional benefits include protecting the skin against harmful UV radiation and facilitating cooling through perspiration.^1–3^ Therefore, understanding the physical and chemical structure of hair fibres is crucial for maintaining its desirable properties and hence it has been the focus of much research for over 150 years.^4^

The hair fibre cross-section consists of three main regions, the medulla, the cortex and the cuticle.^5^ The innermost region, the medulla, comprises vacuolated cells distributed along the centre of the fibre and bound together within a matrix and held by the surrounding cell membrane complex (CMC).^6^ The CMC consists of a central proteinaceous and polysaccharide-rich δ-layer, surrounded by an inner and outer lipid rich β-layer, the most prevalent of which is 18-methyleicosanoic acid (18-MEA).^7–9^ The majority of the hair is made up of the cortex which consists of tightly packed, long cortical cells which are aligned parallel to the hair fibre and embedded within corticular CMC.^3,10^ The outermost region of the hair fibre comprises cuticle layers which serve as a protective outer film consisting of 5 – 10 overlapping scales (cells) of approximately 0.5 to 1.0 µm thickness. Individual cuticle cells have an internal layered structure consisting of the epicuticle, A-layer, exocuticle and endocuticle as shown schematically in Figure 1, and are separated from each other by cuticular CMC.^3,10–13^

**Figure 1:**
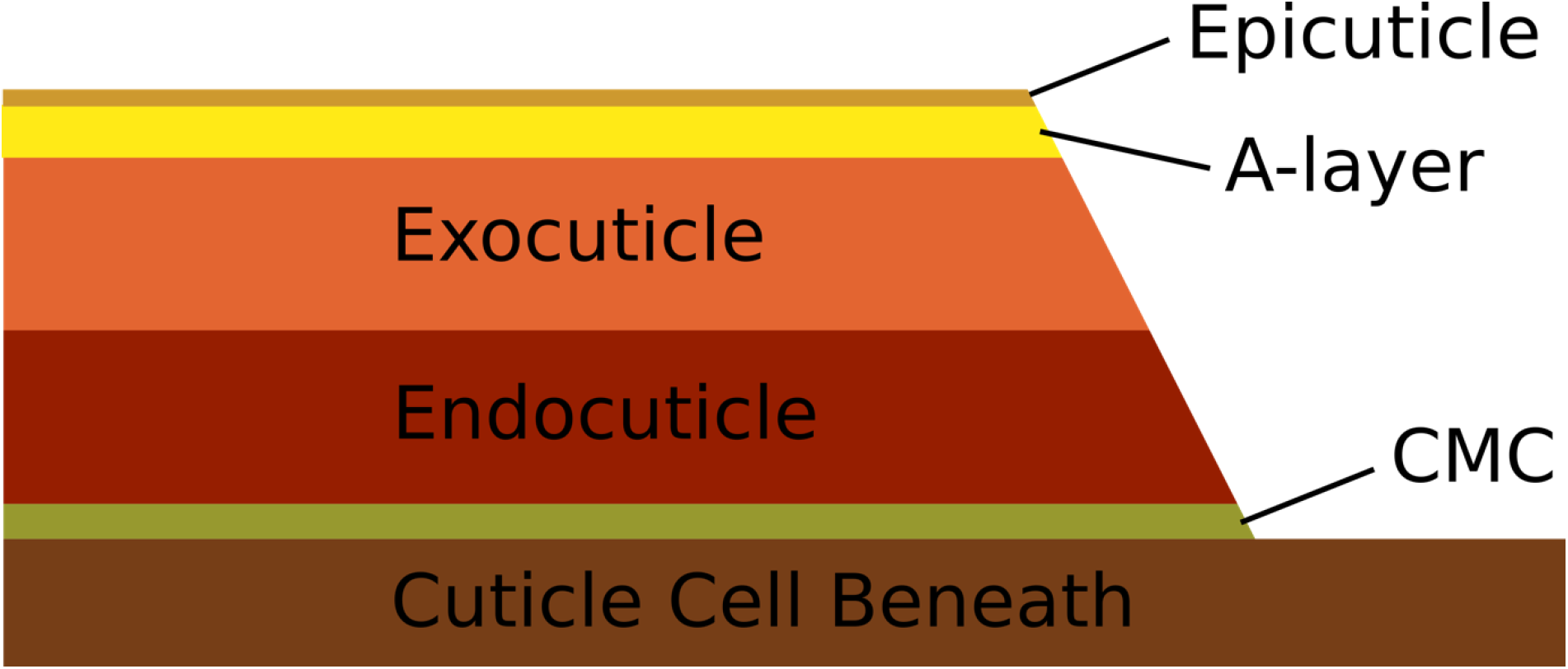
Diagrammatic representation of the layered sub-structure of the cuticle cell.

The outermost layer of the cuticle, the epicuticle, has been shown to be approximately 13 nm thick and to consist mainly of protein (∼80%) with small amounts of lipid and no evidence of carbohydrate.^11,14,15^ In addition, the external surface of the outermost cuticle is covered by a lipid β-layer (18-MEA), originating from the CMC (where the δ- and second β-layer are no longer present).^7^ The layer beneath, the A-layer, provides significant physical and chemical resistance to external stimuli due to its high cystine concentration (the oxidative product of the combination of two cysteine molecules containing a S-S linkage) and hence extensive cross-linking. This layer is therefore high in protein content with only small amounts of lipid.^16,17^ The thickest layer, the exocuticle, lies beneath the A-layer and also has a significant protein network and high cystine concentration. However, at only 15% cystine, the lower concentration compared to the A-layer results in less cross-linking and a smaller, but still significant, physical and chemical resistance.^18^ The final, innermost, layer is the endocuticle which is high in protein but with only 3% cystine and does not contribute much to the resistive properties of the cuticle.^18^

The CMC separating individual cuticle cells binds them together. As mentioned above, the lipid 18-MEA is also the major constituent of the external β-layer, where it covalently binds the S-moieties in the epicuticle and provides the first defensive barrier against external agents.^7–9,19^ This lipid layer has been the focus of much research since it is not only a protective barrier but also affects the physical properties of the external surface of hair like smoothness and hydrophobicity.^20,21^ In general, characterising the physics and chemistry of the cuticle has been of particular importance in research because it provides the fundamental protection and insulation properties of hair.

Previous work using vibrational spectroscopy to study hair structure, such as ATR-FTIR and Raman spectroscopies, provided bulk spectra with some depth profiling of the fibre.^22,23^ Chemical microscopies e.g confocal Raman, have been applied to determine chemical variation through the hair fibre using microtome cross-sections.^24^ These methods yield relatively poor spatial resolution however, owing to the diffraction limit of light, and thus cannot reveal many crucial sub-micron features of the fibre. In contrast, AFM-IR is a hybrid technique that integrates established Atomic Force Microscopy (AFM) with a tuneable infrared (IR) laser to yield spectral mapping of the surface at nanometre resolution. It works by using the AFM tip to detect the localised thermal expansion that follows IR absorption by a vibrational transition of surface molecules and therefore can circumvent the diffraction limit.^25^ In earlier work, Marcott and co-workers used AFM-IR to show localisation of structural lipids in the medulla and cuticular CMC between individual cuticle layers using maps of the ratio of lipid (using 2924 and 2930 cm^−1^ bands corresponding to aliphatic methylene stretching modes) to keratin proteins (2960 cm^−1^ methyl asymmetric stretch, and 1525 cm^−1^ Amide-II band).^26^ The current work aims to characterise the chemical structure of the cuticle itself using the high spatial resolution of AFM-IR, focusing on the Amide-I, lipid carbonyl and cystine-related bands.

## Results

### Cuticle Surface Approaching the Scale-Edge

Figure 2 (a) shows a representative topographical map of the overlapping scale structure of the cuticles on the hair surface. Figure 2(b) is the complementary infrared intensity map at 1630 cm^−1^, corresponding to the Amide-I band, showing that the intensity decreases towards the edge of the cuticle. This arises either because of a lower β-sheet protein secondary structure concentration, that dominates the Amide-I band contribution at ∼1630 cm^−1^, or because the IR laser penetrates less deeply into the cuticle leading to a lower sampling depth. Two further possibilities that could lead to the decreased intensity were considered: inadequate tracking of the contact resonance frequency of the cantilever due to changing mechanical properties of the surface on approaching the edge; and the mechanical differences broadening and decreasing the intensity of the contact resonance peak. These can both be discounted since the observed changes to the contact resonance were small and the PLL appeared to track these efficiently. Additionally, the apparent loss in intensity can also be seen in the spectra of Figure 3 for which the laser is tuned to the contact resonance at each surface location.

**Figure 2:**
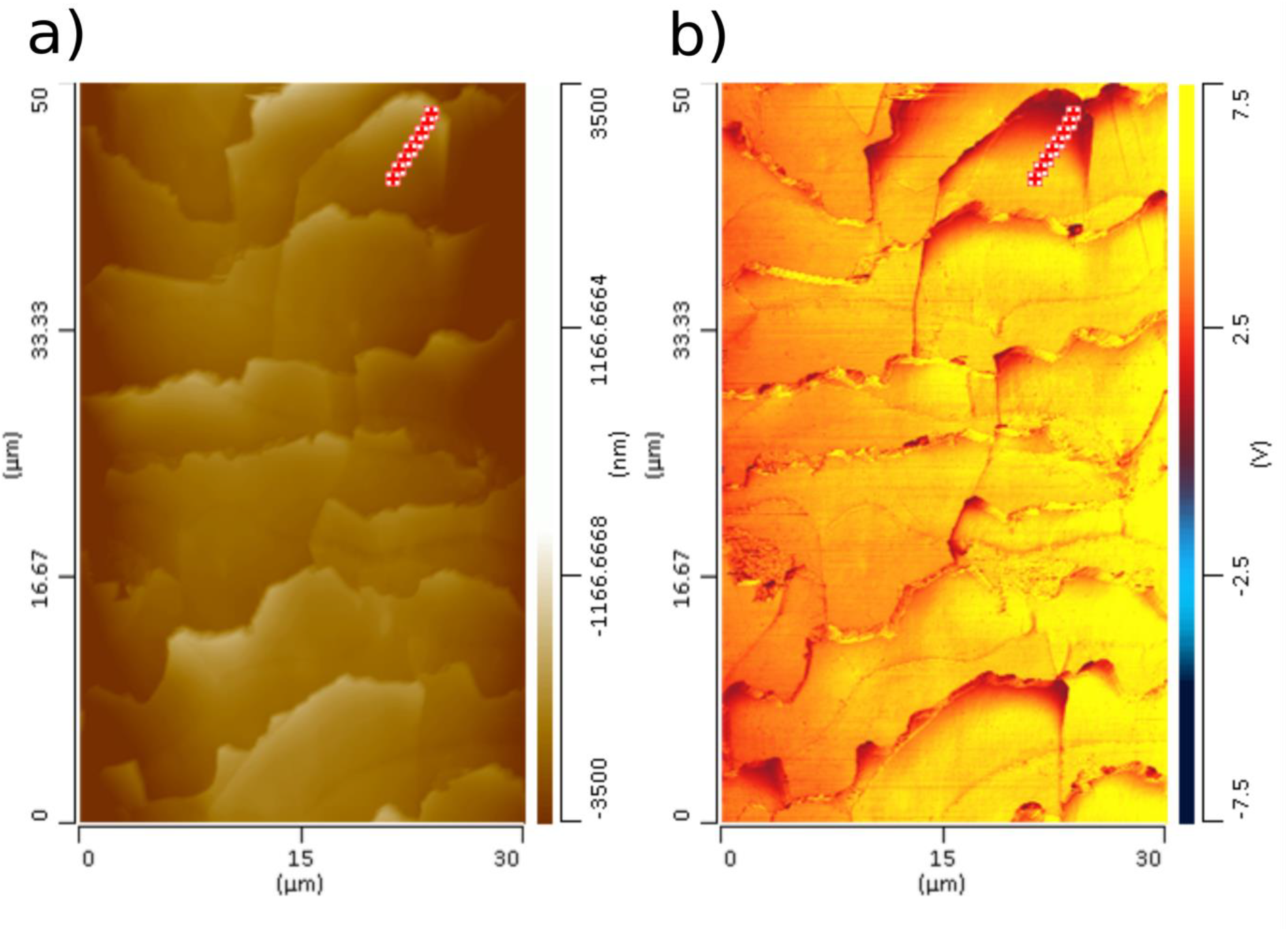
AFM-IR (50 × 30 µm) maps of the external surface of a collection of hair cuticles from an untreated European brown hair showing a) height topography b) IR intensity at 1630 cm^−1^ corresponding to the Amide-I band. The markers in the top right corner of the maps are the positions A to H where spectra were recorded as reported in Figure 3 and Figure 4. Position A is closest to the scale-edge moving incrementally away from the edge to position H.

**Figure 3:**
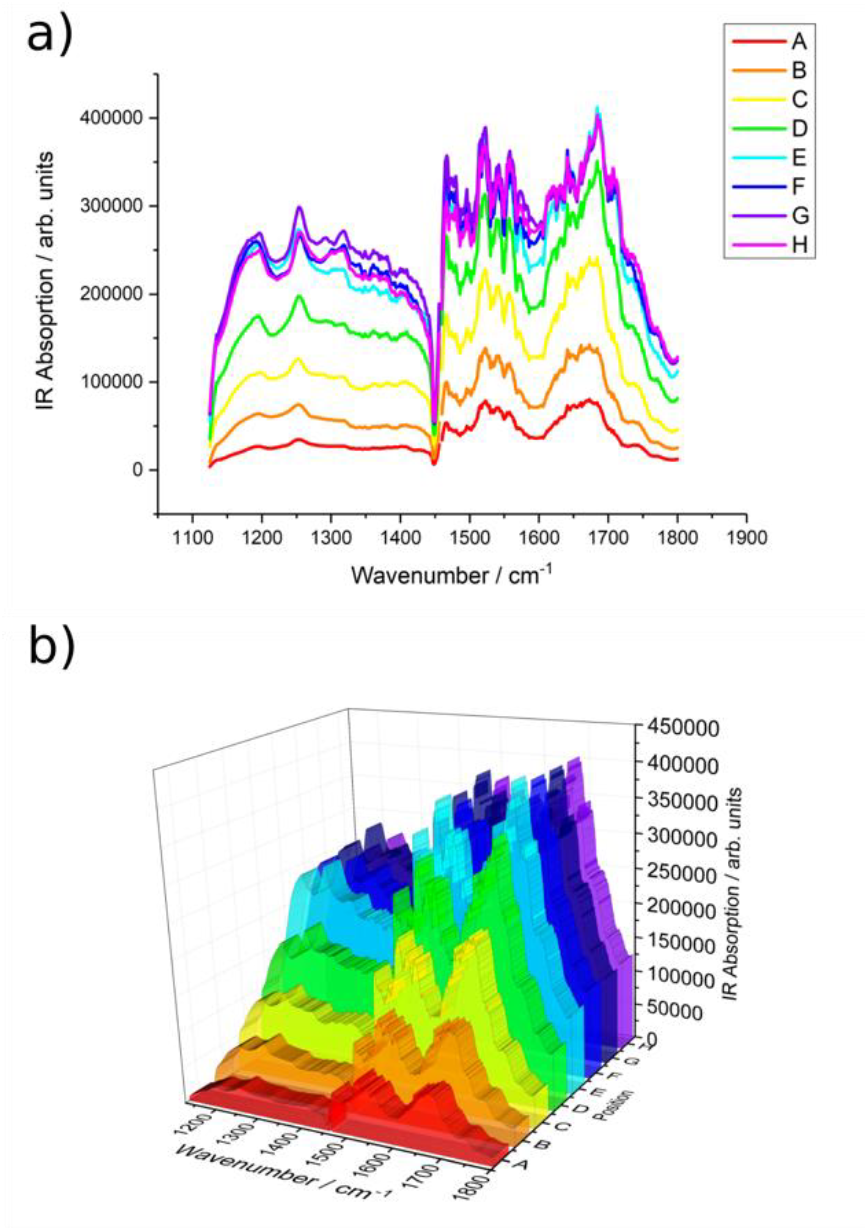
Survey AFM-IR spectra for positions A to H near the edge of a cuticle indicated by the markers in Figure 2. a) line spectra, noise reduced using 20-point smoothing using the Savitsky-Golay algorithm. b) corresponding waterfall plot of the spectra.

To determine the origin of the spectral intensity variation, a series of survey spectra were recorded from the edge of the cuticle cell inwards at the markers indicated in Figure 2. Figure 3 shows the individual spectra (Figure 3(a)) and the corresponding waterfall plot (Figure 3(b)). It can be seen that there is an intensity decrease across the whole spectrum for positions closer to the edge, showing that the drop in intensity at 1630 cm^−1^ nearer the edge is due to an overall decrease in intensity rather than a specific decrease of the Amide-I β-sheet band contribution. This means that, to analyse apparent changes in the protein secondary structure, and chemical composition generally, in proximity to the edge of the cuticle cell, relative proportions of individual vibrational bands to the total IR intensity must be considered. It has been shown that changes in protein secondary structures cause significant variation in the amide bands and in particular in the Amide-I band due to the different environment-dependent contributions of the amide groups in each secondary structure.

Figure 4(a) shows spectra in the Amide-I and lipid carbonyl stretching region for the positions A to H where A is closest to the edge of the cuticle cell. The spectra have been smoothed with a 7-point FFT-filter in order to determine the band contributions using the second-derivatives of the absorption spectra as shown in Figure 4(b).^27^ (Additionally, fourth-derivative spectra were also generated. All the second- and fourth-derivative spectra at each of the points A to H are shown in Supporting Information, S1.) It is clear in Figure 4(b) that each of the spectra have the same contributing bands with minimal shifts in the band origin and hence the significant changes in the spectra in Figure 4(a) are mainly due to the change in relative intensities of the band peaks. Once the band centres had been identified, the IR intensity that each band contributes to the spectra at each position (A to H) from the cuticle cell edge was determined. The band positions and their proportions relative to the total Amide-I intensity of experimentally identified bands in the Amide-I and lipid region are presented in Table 1. The variation in each of the identified contributions, peaks 1 to 9, to the Amide-I and lipid carbonyl region progressing further from the cuticle cell edge are shown in Figure 5. The assignment of each of the peaks, based on band positions and literature assignments, is given in Table 2.^23,28^ Using these assignments, in combination with the upper and lower panels of Figure 5, the molecular contributions at each of the positions A to H can be deduced and are presented in graphical form in Figure 6. These plots show high proportions of α-helices and random coil structures near the edge (position A) and higher proportion of β-sheets and lipid structures further inwards (position H).

**Table 1:** The positions (cm^−1^) and intensities relative to the total Amide-I intensity of the individual bands in the Amide-I and lipid spectral region at sampling positions A to H on top of and near the edge of the cuticle cell.

| Position | A | B | C | D | E | F | G | H |
| --- | --- | --- | --- | --- | --- | --- | --- | --- |
| Peak 1 | 1737<br>0.072 | 1737<br>0.076 | 1737<br>0.080 | 1736<br>0.094 | 1736<br>0.106 | 1735<br>0.113 | 1736<br>0.119 | 1736<br>0.118 |
| Peak 2 | 1723<br>0.073 | 1723<br>0.080 | 1723<br>0.086 | 1722<br>0.102 | 1722<br>0.116 | 1722<br>0.124 | 1722<br>0.128 | 1722<br>0.127 |
| Peak 3 | 1707<br>0.112 | 1708<br>0.116 | 1709<br>0.121 | 1708<br>0.143 | 1709<br>0.157 | 1710<br>0.156 | 1710<br>0.162 | 1709<br>0.162 |
| Peak 4 | 1688<br>0.180 | 1689<br>0.174 | 1689<br>0.181 | 1688<br>0.193 | 1688<br>0.195 | 1689<br>0.187 | 1689<br>0.189 | 1689<br>0.187 |
| Peak 5 | 1670<br>0.200 | 1674<br>0.191 | 1673<br>0.192 | 1672<br>0.185 | 1672<br>0.183 | 1672<br>0.177 | 1672<br>0.177 | 1673<br>0.179 |
| Peak 6 | 1656<br>0.187 | 1661<br>0.188 | 1663<br>0.184 | 1663<br>0.175 | 1662<br>0.167 | 1662<br>0.165 | 1663<br>0.167 | 1663<br>0.167 |
| Peak 7 | 1644<br>0.180 | 1643<br>0.180 | 1644<br>0.175 | 1645<br>0.168 | 1645<br>0.166 | 1644<br>0.166 | 1643<br>0.163 | 1644<br>0.163 |
| Peak 8 | 1625<br>0.139 | 1626<br>0.149 | 1626<br>0.145 | 1626<br>0.147 | 1627<br>0.147 | 1626<br>0.154 | 1625<br>0.152 | 1628<br>0.154 |
| Peak 9 | 1613<br>0.113 | 1612<br>0.118 | 1613<br>0.123 | 1614<br>0.133 | 1615<br>0.142 | 1615<br>0.151 | 1614<br>0.151 | 1615<br>0.152 |

**Table 2:** Approximate band origins and their assignments in the Amide-I and lipid spectral region for positions A to H.

| Peak | Approximate Peak Centre / $\text{cm}^{-1}$ | Assignment |
| --- | --- | --- |
| 1 | 1736 | Lipid |
| 2 | 1722 | Asp / Glu Acidic Side-Chains |
| 3 | 1709 | Asp / Glu Acidic Side-Chains |
| 4 | 1689 | Asn / Gln Amide Side-Chains |
| 5 | 1672 | β-Turn |
| 6 | 1662 | Random Coil |
| 7 | 1644 | α-Helix |
| 8 | 1626 | β-Sheet |
| 9 | 1614 | β-Sheet |

**Figure 4:**
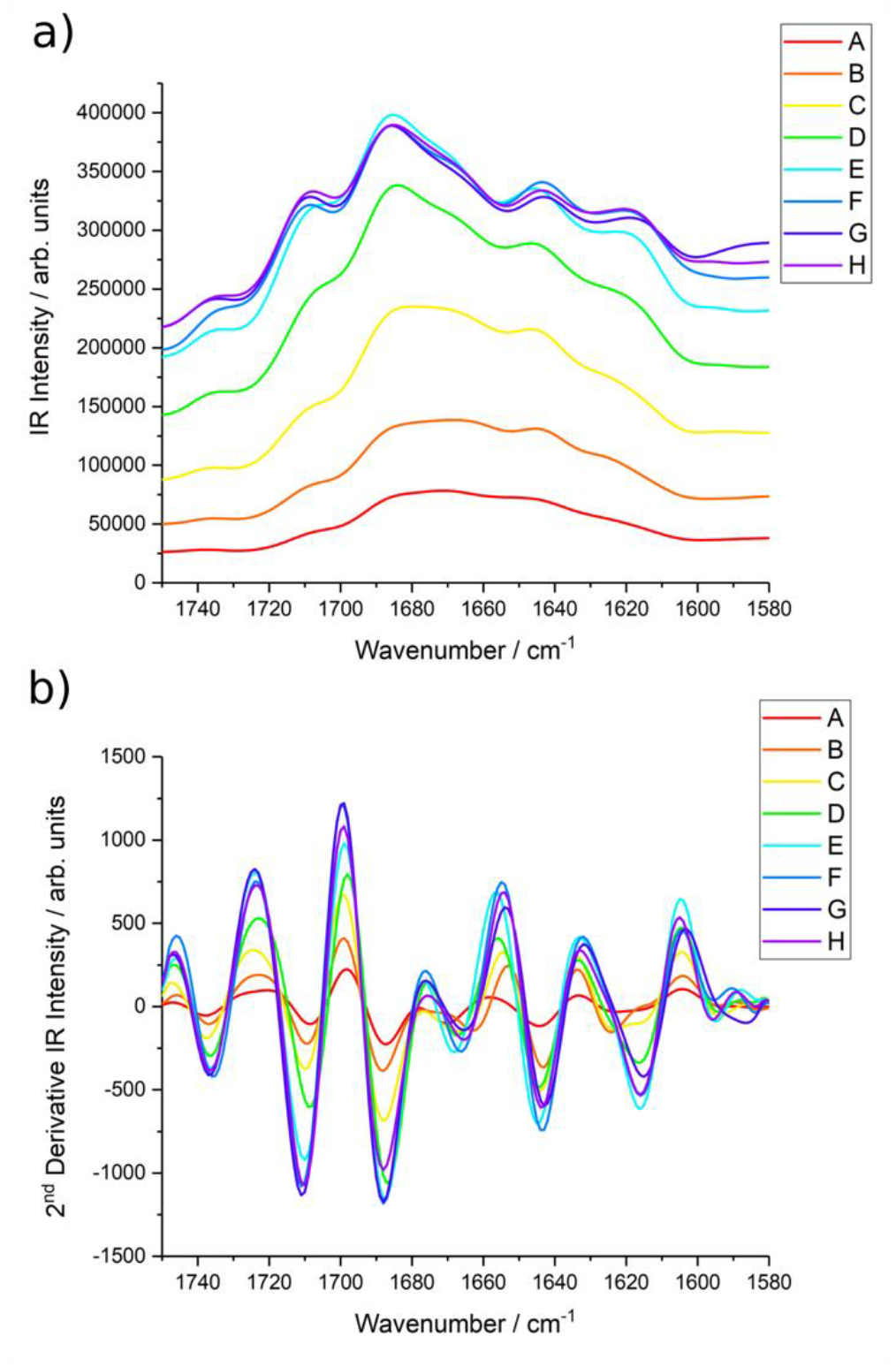
AFM-IR spectra in the region of the Amide-I band at positions A to H. a) recorded spectra smoothed using a 7-point FFT-filter to reduce noise. b) Second-derivative spectra.

**Figure 5:**
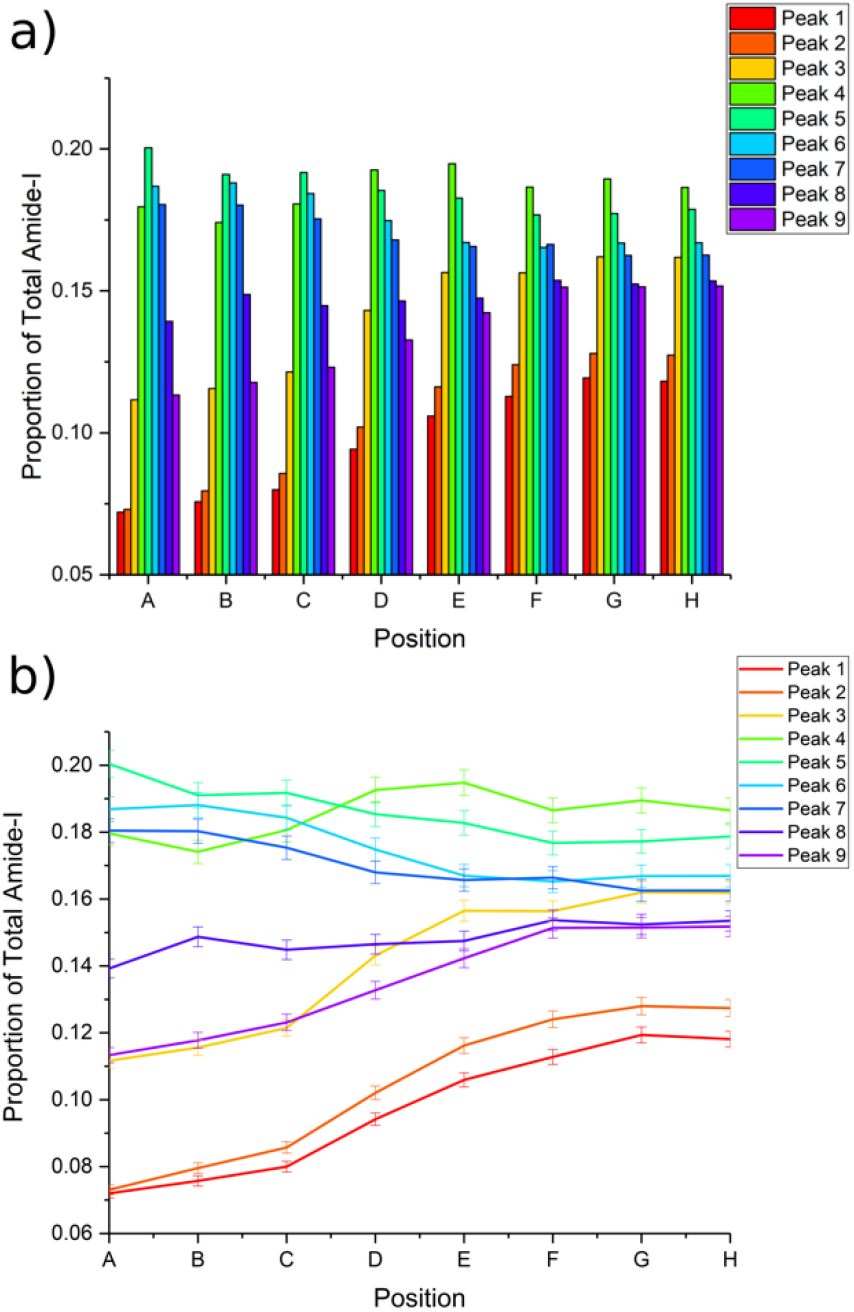
a) column plot showing the variation in intensity of the nine identified bands as a proportion of the total Amide-I intensity at each of the sampling points A to H. b) line plot of individual bands as a proportion of the total Amide-I intensity as a function of the sampling points A to H.

**Figure 6:**
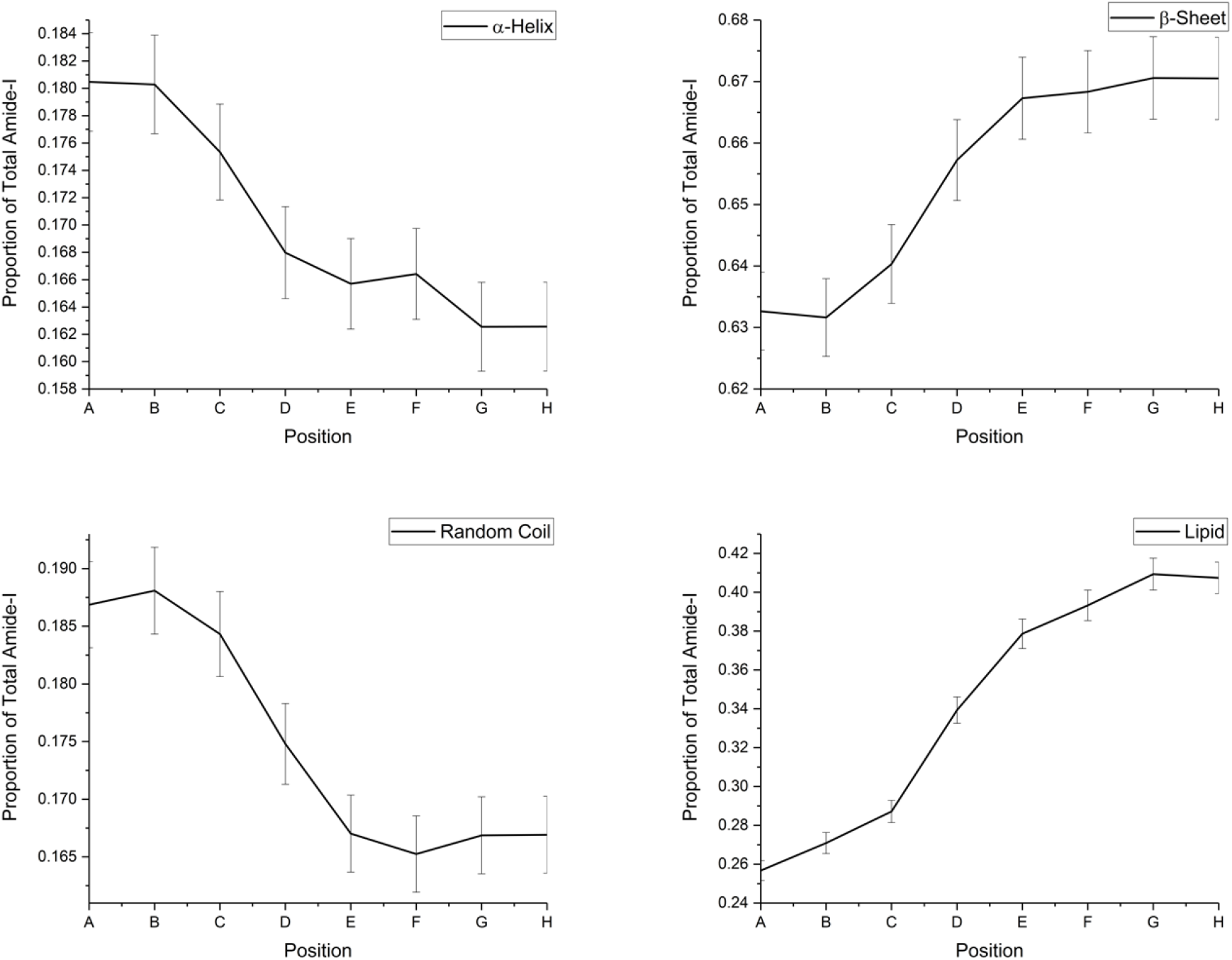
Variation in protein secondary structures and lipid as a proportion of the amount of total Amide-I determined from the total band intensity contribution to the Amide-I and lipid regions for positions A to H.

The structural features of different protein secondary structures are well known. Specifically, the α-helix consists of an un-stretched rod-like coil and, other than in tensile deformation, imparts increased rigidity and stiffness to protein structures.^29,30^ β-sheets, on the other hand, consist of stretched chains hydrogen-bonded into extensive sheets that impart greater flexibility and lower stiffness than the α-helices.^29,30^ The data in Figure 6 shows that the cuticle edge has a significant α-helix component and hence greater stiffness than positions further from the edge. This is in agreement with expectation since the edges require a greater stiffness and rigidity. The β-sheet structure is the natural stable protein conformation and also has been shown to result from applied stress whereby coils are unravelled into extended chains which subsequently cross-link via H-bonds into the extended sheet structures.^29,30^ The edge of the cuticle cells show greater proportions of α-helices and random coil structures, whilst towards the centre of the cuticle cell β-sheets are more prevalent, as shown in Figure 6. This could suggest that the edge regions are under less stress indicating that the scales are generally under tension. However, it is also possible that the proteins in the cuticle cells are naturally β-sheet structures due to the fundamental stability of this structure rather than arising from conversion of α-helices due to external stress experienced during the fibre growth phase. It is worth noting that, in all the spectra, the β-sheet contribution is the major component which supports the conclusion that the scales generally are under tension as well as that the proteins exist in their naturally stable conformation. Further work is required to elucidate the origin of these structures.

The survey spectra in Figure 3 show an overall decrease in intensity near the cuticle cell edge which is probably due to lower IR penetration because of the air gap below the cuticle cells, which is greatest at the edge. This means that spectra at the edge are weaker because they arise from the outermost part of the cuticle whereas further in there is the possibility of greater penetration of the IR to lower layers i.e. deeper cuticle cells and even the cortex. Although the laser intensity may be sufficient to excite vibrations deeper into the hair, the resulting thermal expansion on absorption will dissipate more significantly in deeper layers than closer to the surface. Clearly this effect enhances the surface selectivity of the recorded spectra. Nevertheless, although it may be a diminished contribution to the spectrum, it is important to consider whether variations in protein structure come exclusively from the cuticle cells themselves. The underlying cortex, in its coiled fibrillar state, is generally under fairly low stress and consists predominantly of an α-helix structure. Hence, if the spectra of the cuticle cell (in regions away from the edge) had a significant contribution from the underlying cortex they should show a greater presence of α-helix bands which is not what is observed experimentally (Figure 6). It is, therefore, concluded that the AFM-IR technique is surface-selective and that contributions to the spectra from below the cuticle are insignificant.

### Cuticle Cell Layered Sub-Structure

As well as studying the cuticle surface, we have used AFM-IR to examine the edge of the cuticle cell, the schematic structure of which was shown earlier in Figure 1. Figure 7 comprises the AFM-IR results for the edge of the cuticle cell, showing topographical maps ((a) and (c)) and IR intensity maps at 1650 cm^−1^ ((b)) and 1730 cm^−1^ ((d)), corresponding to the Amide-I band from α-helix protein secondary structures and lipid carbonyl stretching bands respectively. Although the layered structure is poorly resolved in the topography, it is sharply resolved in the IR intensity maps, indicating that there is significant variation in the chemical structure and, therefore, in the composition of the cuticle cell layers. This experimental layering observed by AFM-IR correlates well with the schematic view in Figure 1.

**Figure 7:**
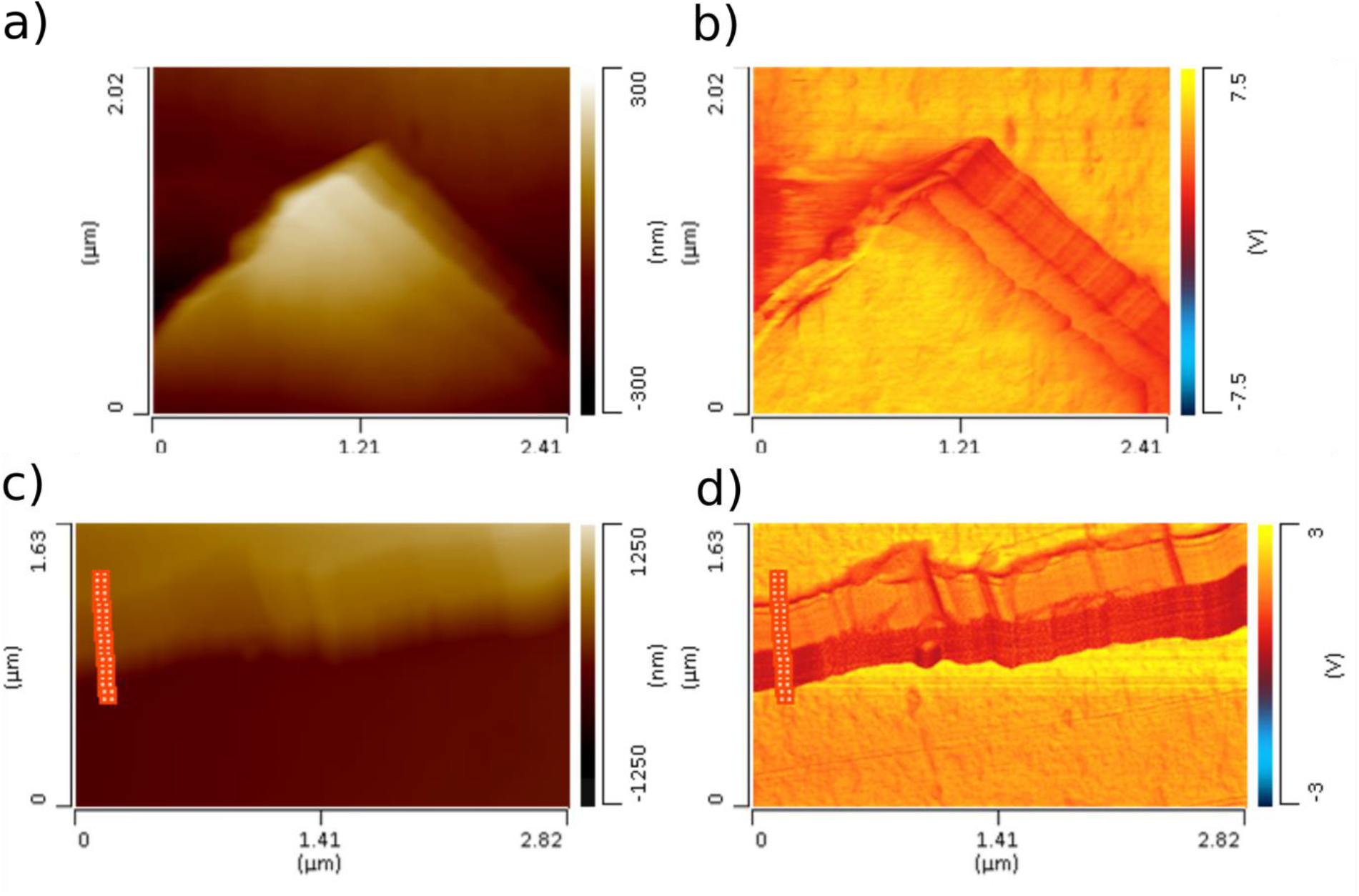
Maps of the height (left panels: (a) and (c)) and IR intensity (right panels: (b) and (d)) of the cuticle cell edge recorded by AFM-IR. The IR intensity maps were recorded at (b) 1650 cm^−1^ and (d) 1730 cm^−1^. Markers in (c) and (d) correspond to the positions 1 to 20 where spectra were recorded (starting from the top of the image and progressing downwards).

Next, a spectral line-array was recorded across the edge of the cuticle cell at the points indicated by the markers in Figure 7, (c) and (d). The individual spectra, labelled subsequently from 1 to 20, commence at the outermost cell surface of the cuticle and end on the external surface of the cell beneath (i.e. traversing the edge). The recorded spectra and their second derivatives are shown in Figure 8. These spectra have been smoothed in an analogous manner to those in Figure 4 to yield accurate band centres from the second and fourth derivatives of the recorded spectra. The band origins and their corresponding intensities are presented in Table S2. This data was used to generate a graphical presentation of the Amide-I composition at each sampling point (Figure 9(a)) and as well as the variation of individual band contributions across the 1 to 20 sampling positions (Figure 9(b)). Linking the observed layers to the positions where individual spectra were recorded (positions 1-20) yields a close correlation between changes in the spectral fingerprint and the observed layered structure of the cuticle cell.

**Figure 8:**
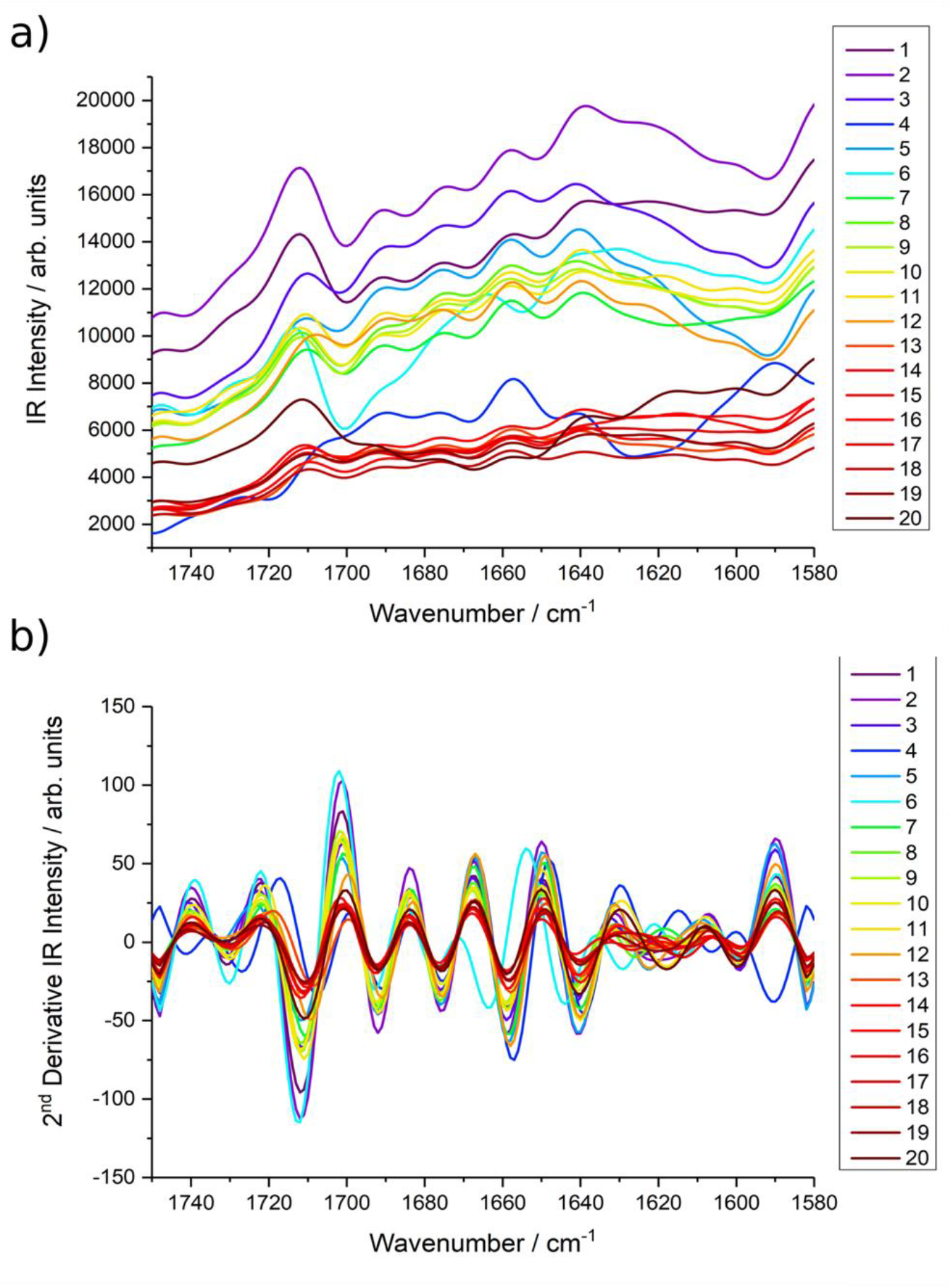
AFM-IR spectra in the region of the Amide-I band at positions 1 to 20. a) recorded spectra smoothed using a 7-point FFT-filter to reduce noise. b) Second derivative spectra.

**Figure 9:**
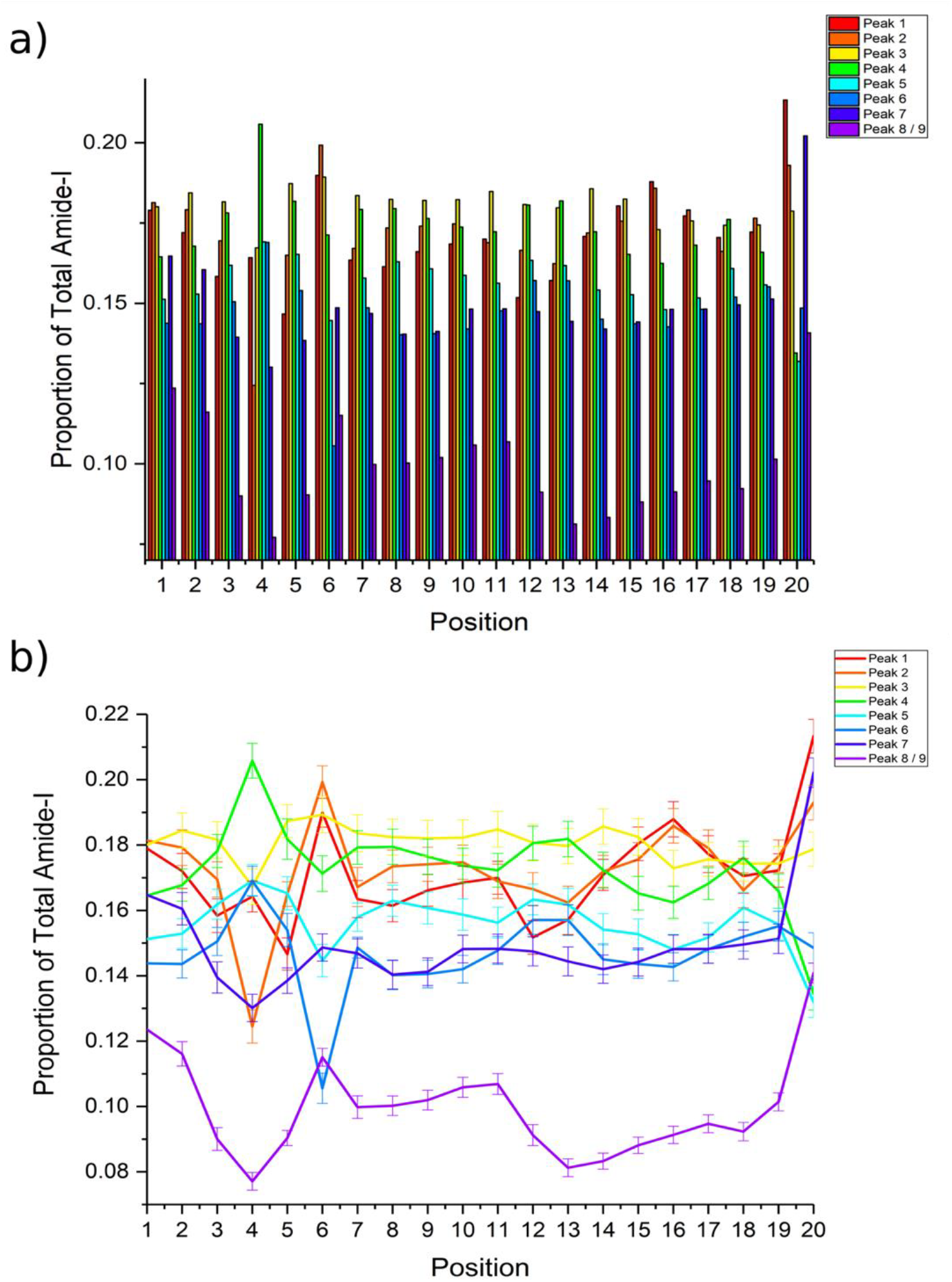
a) column plot showing the variation in intensity of the nine identified bands as a proportion of the total Amide-I intensity at each of the sampling points 1 to 20. b) line plot of individual bands as a proportion of the total Amide-I intensity as a function of the sampling points 1 to 20. Uncertainties were calculated from the relative proportion of the spectral noise level.

It is instructive to examine the ratio of the amide to lipid contributions and these can be deduced from the spectral line-array shown in Figure 7(c) and (d). Figure 10 shows the ratio of the protein (Amide-I, bands 1-6) to the lipid (most significant carbonyl C=O stretching mode, bands 7-9) contributions at each of the sampled positions. Initially, there is a lower protein contribution, and conversely a high lipid concentration, rising to a maximum at position 4. This subsequently drops away remaining roughly constant until position 11 after which it increases again up to position 14 until finally it starts to decrease again until suddenly dropping at position 20. Although these changes indicate that there are differences in chemical composition through the sample it is difficult to quantify the amounts in different layers because of the unknown exact penetration depth of the IR laser. Nevertheless, the observed changes are consistent with what is known about the different cuticle cell layers.

**Figure 10:**
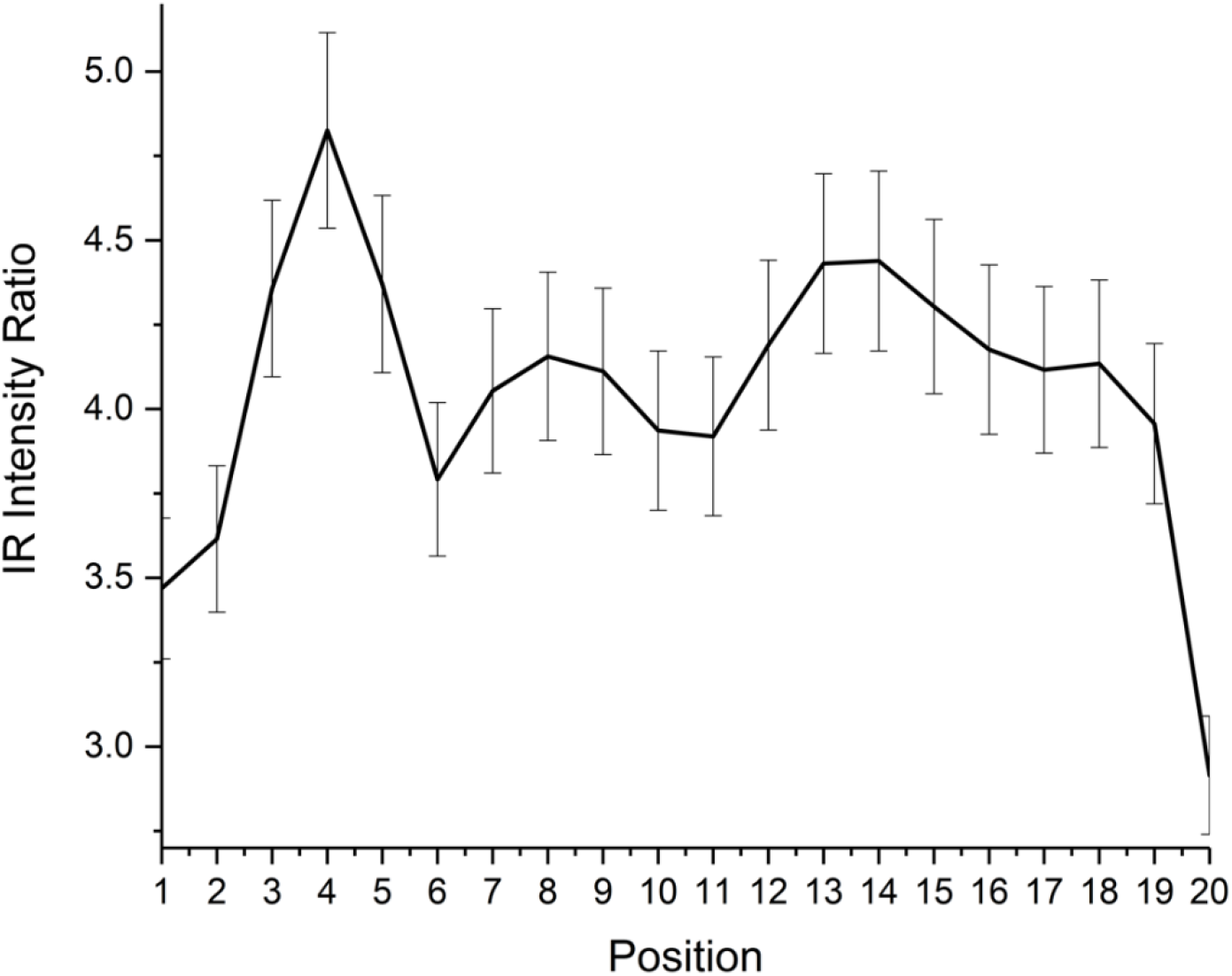
The protein:lipid IR intensity ratio calculated from the Amide-I and lipid carbonyl bands at each of the sampling positions 1 to 20.

Initially the AFM tip probes the outer β-layer that is high in lipid content (18-MEA) giving the low protein-to-lipid ratios found for positions 1 and 2. The higher ratio at positions 3 to 5 corresponds to the higher protein content of the epicuticle and underlying A-layer. The exocuticle begins at position 6, where the protein content drops. This lower protein content lasts until position 12/13 which corresponds to the clear boundary between the two thick layers. This is apparent in the IR intensity map in Figure 7 which marks the transition from the exocuticle to the endocuticle. After this boundary, the protein content rises. From position 14-19, the ratio appears to fall slowly until it abruptly drops at position 20 as the tip probes the β-layer of the next cuticle cell. The gradual drop in protein content after position 14 is likely due to IR penetration through the endocuticle leading to intensity contributions from the CMC underneath, which is comparatively high in lipid content.^7–9^ Furthermore, some lipid from the CMC can potentially penetrate into the endocuticle, decreasing in concentration the further it gets from its origin in the CMC but nevertheless affecting the protein-to-lipid ratio.^31^ However, this observed drop falls within the uncertainty margins hence any explanation of the cause of the ratio change cannot be supported.

As well as the variable amounts of protein and lipid in the cuticle layers there is a large difference in the cystine content of the layers which affects the amount of cross-linking of the protein networks. In order to gain insight into the role of cystine and more specifically cystine-related products in the cuticle cell structure, AFM-IR spectra between 1125 and 1350 cm^−1^ were recorded. This region also contains the Amide-III bands. Savitzky-Golay-smoothed spectra for positions 1 to 20 across the cuticle cell edge are presented in Figure 11. Although the spectra contain fewer obvious features, second and fourth derivative processing using FFT-smoothed spectra clearly identifies several contributing peaks with the band origins given in Table 3. Because the cuticle cells make up the surface of the hair, they are susceptible to oxidation. When this occurs, it converts the cystine into oxidation products like cystine monoxide, cystine dioxide, cysteine-S-thiosulphate and cysteic acid with progressive oxidation eventually leading to cleavage of the cystine S-S bond.^32^ Some of the bands in Table 3 have been assigned to these oxidation products and provide a measure of the total cystine content.^23,33–37^

**Figure 11:**
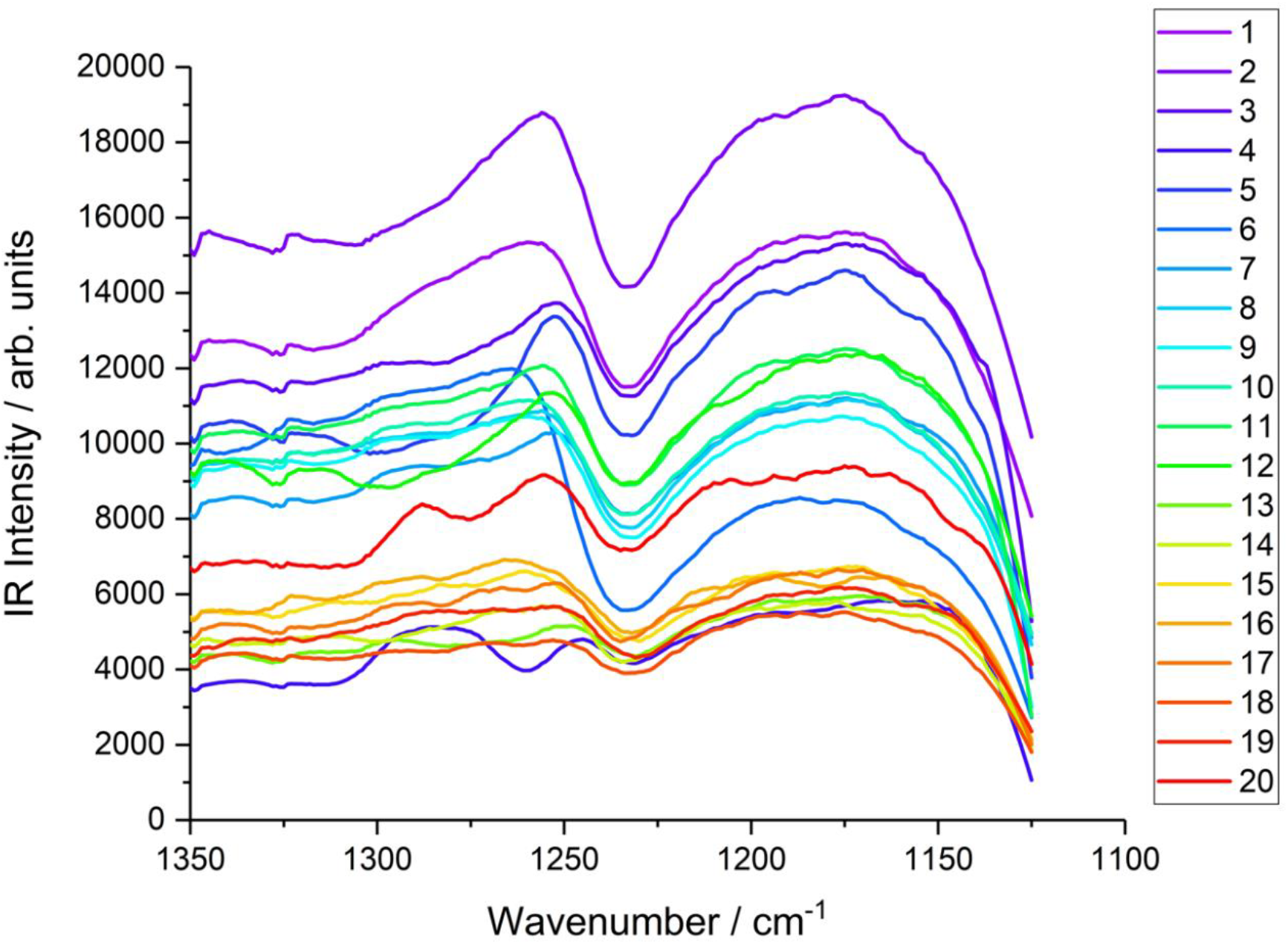
AFM-IR spectra in the Amide-III and cystine-related compound region 1125 – 1350 cm^−1^ at positions 1 to 20. Spectra have been smoothed using a 20-point Savitzky-Golay algorithm.

**Table 3:** Approximate band origins and their assignments in the cystine and Amide-III spectral region.

| Peak | Approximate Peak Centre / $\text{cm}^{-1}$ | Assignment |
| --- | --- | --- |
| 1 | 1137 | SO<sub>2</sub> stretch |
| 2 | 1154 | C-N stretch |
| 3 | 1170 | Asymmetric Cysteic acid |
| 4 | 1186 | SO<sub>2</sub> (sulfonyl chloride) |
| 5 | 1201 | Cysteine-S-sulfate |
| 6 | 1210 | S-O stretch / Amide-III |
| 7 | 1215 | S-O stretch / Amide-III |
| 8+ | >1225 | Various Amide-III Secondary Structures |

The ratio of selected cystine-related bands to selected Amide-III bands are plotted as a function of positions 1 to 20 in Supporting information, S3. Bands 1 and 3 in Table 3 were selected as input for cystine data, chosen as the most prominent bands assigned in earlier work on hair spectra, and bands at 1228 and 1242 cm^−1^ for Amide-III data, because they do not coincide with cystine-related compounds or other features in this spectral region. The most striking feature in Figure S3 is the significant drop of nearly 10% in the ratio between positions 13 and 18, the region of the endocuticle (clearly identifiable in the IR intensity map in Figure 7(d)). This provides unambiguous confirmation of literature reports that the A-layer and exocuticle are relatively cystine-rich while the endocuticle is cystine-poor. A less prominent cystine-poor region occurs at positions 1 and 2 which agrees with the known composition of the epicuticle and outer β-layer. Interestingly, the apparent increase in cystine content at positions 18 and 19 is actually believed to be due to a contribution from the glycosidic link stretching band in the polysaccharide-rich δ-layer (or similar C-O stretch from triglycerides in this layer) in the underlying CMC.^38–41^ This increase drops at position 20 since the position 20 spectrum is from the top of another cuticle cell where only the β-layer of the CMC is present.

The significance of the analysis of the data presented in Figure S3 is reduced by the uncertainties indicated by the error bars on the plot which are larger than those in the analogous plots in Figure 6 and Figure 10. These uncertainties arise mainly from the noise level in the spectra which therefore results in variations in relative peak ratios. The only clear variation in the cystine-to-protein ratio, lying outside the limits imposed by the uncertainties, is the drop in cystine observed between positions 13 and 18 as well as the subsequent increase for positions 18-20 as discussed above. Therefore, to reduce the uncertainties, rather than utilising specific band contributions, the total band areas were measured by peak-integration to give the cystine to protein ratio. Despite the band between 1125-1225 cm^−1^ containing some contributions of protein origin, the band above 1225 cm^−1^ in this region contains no cystine-related contributions hence the ratio of the two integrated areas gives an alternative albeit qualitative indication of the relative proportions of protein and cystine. This ratio is shown in Figure 12 where a clear reduction in the magnitude of the uncertainties can be observed. Furthermore, there is a similar ratio profile to that in Figure S3. This further emphasises the high cystine concentration in the A-layer (shown here to be even greater than in the exocuticle which couldn’t be distinguished in Figure S3) with moderate levels of cystine in the exocuticle and subsequent drop for the endocuticle as expected from the known structures of these layers. Again, there is an observed increase in this ratio for position 18 and 19 which are believed to source from alternative vibrations in the CMC as described above. Finally, the drop at position 20 is observed where the tip is now atop the exposed surface of the cuticle cell beneath and so the ratio here is equivalent to that of position 1 as anticipated.

**Figure 12:**
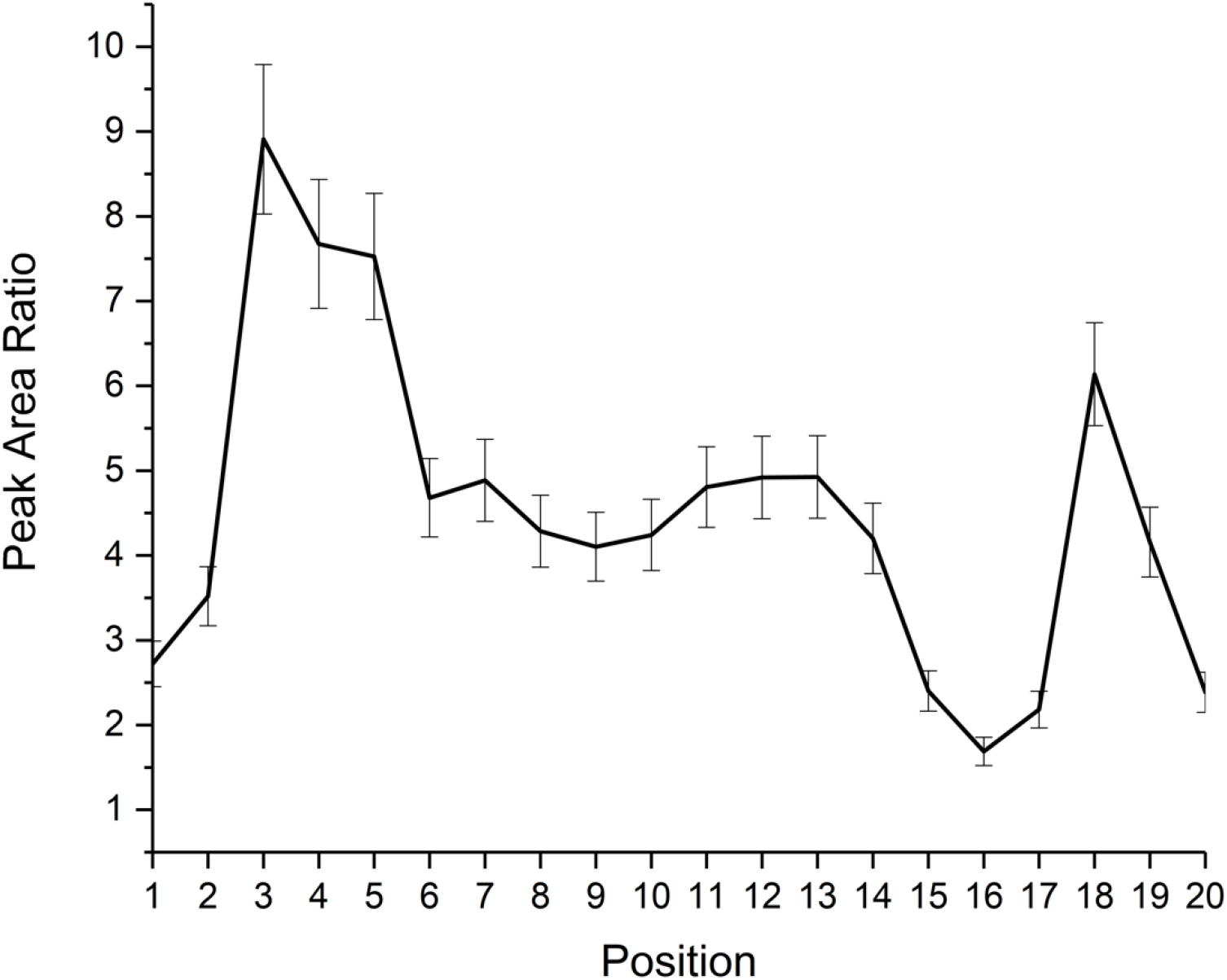
The cystine-to-protein intensity ratio calculated from the sulfur-oxygen and Amide-III infrared band intensities as a function of sampling position 1 to 20.

## Discussion

The layered cuticle cell structure of the external surface of hair has been widely studied but the current work is the first report of surface-selective infrared spectroscopy at nano-scale resolution. Two spectroscopically important constituents have been investigated yielding fundamental information on the distribution of protein and lipid at the surface. Survey measurements of the cuticle cells using AFM-IR showed significant changes to the Amide-I and lipid carbonyl stretching bands on approaching the cell-edge. The spectra show that there is a shift from α-helix and random coil protein secondary structures to β-sheets, although β-sheets remain the prominent structure throughout. β-sheets are known to result from unravelled coils which are under high tension.^29,30^ It is considered that the cuticle cell towards its edges could be under less tension which would explain the lower proportion of β-sheets. In contrast to the proportion of β-sheets, the proportion of α-helices rises towards the edges. α-helices are known to provide greater mechanical rigidity^29,30^ so their increasing proportion on approaching the edges provides the necessary greater rigidity in these more exposed areas. If there is less tension in the edge region, then random coils would also have a higher proportion there, as found experimentally. An alternative explanation for the differences is that the natural stable protein conformation is the β-sheet structure which dominates from the growth process allowing the more stable structure to form. The change to α-helices near the edge therefore results from environmental stresses associated with the edge making the more rigid structure the favoured and resulting conformation.

The lipid content is also observed to be lower at the edges although the three lipid bands detected retain the same relative intensities. The main source of the lipid intensity must be from the external β-layer on top of the epicuticle since the layers below are mainly protein until the CMC is reached at the base of the cuticle cell. As explained earlier, this will only have a weak signal due to the attenuation of the IR intensity by the upper layers as well as the dissipation of the thermal expansion deeper inside the hair. The maintenance of the lipid relative band intensities across the surface means that the proportion of lipid is reduced near the edges and is not due to a change in lipid composition or structure. This is consistent with the stripping of the β-layer which commonly occurs in damaged hair.^20,21,42–44^

The partial splaying that occurs at the edge of the cuticle cell^45^, as well as external damage, as mentioned above with regard to the lipid β-layer, provided the opportunity to investigate the layering of the edge of the cuticle cell. Spectra across the edge, covering individual layers, showed significant changes to the amide, lipid and cystine-related regions. The changes in the spectra revealed a clear layered structure and the interpretation of the spectra confirmed the proportions of lipid to protein as well as the relative cystine content of the different layers. Specifically, the external surface of the β-layer and epicuticle showed a high lipid and lower protein concentration as well as a lower proportion of cystine. In contrast, the next layer, the A-layer, is higher in protein and lower in lipid along with having a higher cystine component. This agrees with the current view of the molecular composition of these layers.^7,11,14–18^ Lower down from the cuticle cell surface, the AFM-IR spectra provided data on the composition of the exocuticle which showed a high cystine content, although not as large as the A-layer, as well as a slightly increased proportion of lipid. The final layer, the endocuticle, showed a high proportion of protein accompanied by a significant reduction in cystine content. It is proposed that the apparent increase in the cystine content at the bottom of the cuticle is the result of penetration of the IR light into the CMC beneath. In addition to the spectral investigations of the layers, the physical thicknesses of the observed layers are in accord with other measurements of, for example, the thickest layers, the endocuticle and the exocuticle, as well as the thinner A-layer and epicuticle at the surface.

## Conclusion

The present study has used AFM-IR to investigate spectroscopic aspects of hair cuticle cells, namely the protein and lipid distribution on the cell surface and the protein and cystine composition of the different layers exposed at the cell-edge. As well as investigating changes in the protein and lipid content by analysing their bands between 1580 and 1640 cm^−1^, cystine containing bands and Amide-III bands lying between 1125 and 1250 cm^−1^ were also investigated. The composition of the cuticle cell surface showed a shift from higher proportions of α-helices and random coil secondary protein structures at the edge to β-sheets further inwards. Additionally, the lipid composition of the surface also increased on moving away from the edge. The different chemical composition found for the layered edge of the cuticle cell is compatible with the known cross section of the cuticle cell layers. Initially the tip probes the outer β-layer which is high in lipid content and low in protein. Below this is the epicuticle and A-layer which are found to have a higher protein content. The protein content drops as the tip probes the next layer, the exocuticle, and finally rises again at the endocuticle, the second of the two thick layers. It was confirmed that the exocuticle layer and A-layer are cystine-rich and the endocuticle layer cystine-poor.

## Methods

Human hairs (European brown) were washed several times with Millipore water and, following drying, mounted on steel substrates with double-sided tape for AFM-IR analysis of the cuticle cells. Spectra and mapping was carried out in contact mode using an Anasys NanoIR2 instrument equipped with a MIRcat Laser system (Daylight Solutions) containing four quantum cascade lasers (QCLs). Recorded IR spectra were smoothed using either the Savitzky – Golay algorithm with 20 points at 1 cm^−1^ resolution, which reduced noise and produced the IR intensities used in quantitative analysis, or a 7-point FFT-filter algorithm for deconvolution to determine band centres from the second- and fourth-derivatives of the smoothed spectra.

## Author Contributions

MTLC designed the experiments and supervised the acquisition of the data.

APF conducted the experiments, processed the results and drafted the manuscript.

PBD oversaw the structure of the manuscript and supervised and refined the interpretation of the data.

## Acknowledgments

We are grateful to Dr Paul Pudney and Dr Ken Lee at Unilever R&D for providing support and guidance through the project. Additionally, we are thankful for Unilever R&D and the EPSRC for providing the iCASE studentship for A. P. F. on Grant EP/R511870/1.

## Supplementary Information

### S1. Second- and Fourth-Derivative Peak Finding Spectra

**S1:**
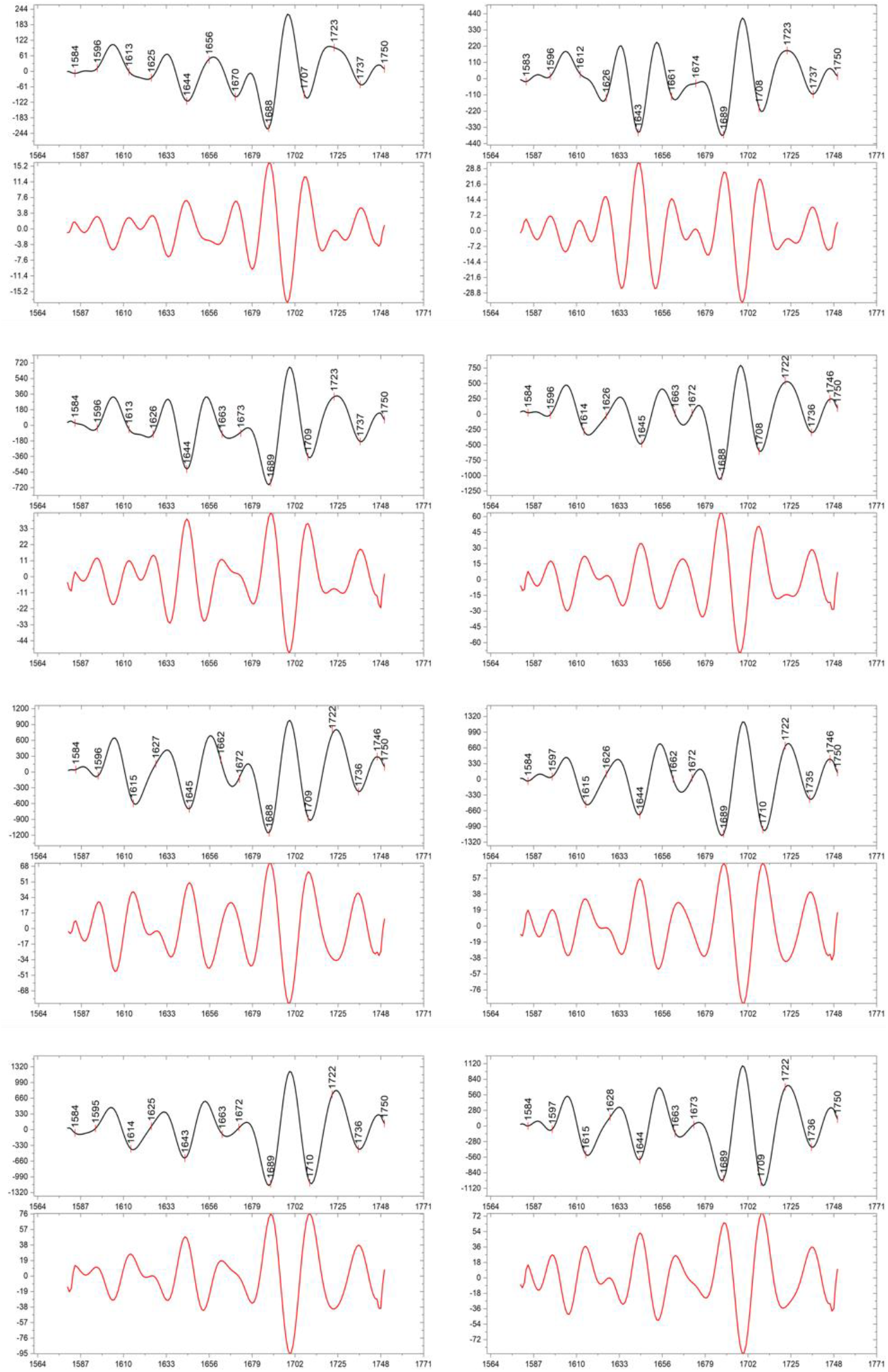
Spectra showing the second- and fourth-derivatives for positions A-H given by the markers in Figure 2 (full spectra shown in Figure 3 and Amide-I and lipid region used to calculate the derivatives are shown in Figure 4).

### S2. The positions (cm^−1^) and intensity contribution to the total Amide-I intensity of individual bands at positions 1 to 20

**S2:** The positions (cm^−1^) and intensity contribution to the total Amide-I intensity of the individual bands in the Amide-I and lipid spectral region at sampling positions 1 to 20 at the edge of the cuticle cell.

| Position | Peak 1 | Peak 2 | Peak 3 | Peak 4 | Peak 5 | Peak 6 | Peak 7 | Peak 8/9 |
| --- | --- | --- | --- | --- | --- | --- | --- | --- |
| 1 | 1614<br>0.179 | 1625<br>0.181 | 1641<br>0.180 | 1659<br>0.165 | 1676<br>0.151 | 1692<br>0.144 | 1711<br>0.165 | 1731<br>0.124 |
| 2 | 1614<br>0.172 | 1625<br>0.179 | 1641<br>0.184 | 1659<br>0.168 | 1676<br>0.153 | 1692<br>0.144 | 1711<br>0.161 | 1731<br>0.116 |
| 3 | 1614<br>0.158 | 1626<br>0.169 | 1642<br>0.182 | 1659<br>0.178 | 1676<br>0.162 | 1692<br>0.151 | 1711<br>0.139 | 1732<br>0.090 |
| 4 | 1607<br>0.164 | 1623<br>0.124 | 1639<br>0.167 | 1657<br>0.206 | 1675<br>0.169 | 1691<br>0.169 | 1708<br>0.130 | 1729<br>0.077 |
| 5 | 1614<br>0.147 | 1624<br>0.165 | 1641<br>0.187 | 1658<br>0.182 | 1675<br>0.165 | 1692<br>0.154 | 1711<br>0.138 | 1734<br>0.090 |
| 6 | 1614<br>0.190 | 1629<br>0.199 | 1645<br>0.189 | 1663<br>0.171 | 1678<br>0.145 | 1694<br>0.106 | 1712<br>0.149 | 1730<br>0.115 |
| 7 | 1612<br>0.163 | 1625<br>0.167 | 1641<br>0.184 | 1658<br>0.179 | 1675<br>0.158 | 1692<br>0.149 | 1710<br>0.147 | 1729<br>0.100 |
| 8 | 1613<br>0.161 | 1626<br>0.173 | 1642<br>0.182 | 1659<br>0.179 | 1676<br>0.163 | 1693<br>0.140 | 1711<br>0.140 | 1730<br>0.100 |
| 9 | 1614<br>0.166 | 1626<br>0.174 | 1642<br>0.182 | 1659<br>0.176 | 1676<br>0.161 | 1693<br>0.141 | 1711<br>0.141 | 1730<br>0.102 |
| 10 | 1613<br>0.169 | 1626<br>0.175 | 1641<br>0.182 | 1659<br>0.174 | 1676<br>0.159 | 1693<br>0.142 | 1711<br>0.148 | 1730<br>0.106 |
| 11 | 1617<br>0.170 | 1625<br>0.169 | 1641<br>0.185 | 1659<br>0.172 | 1676<br>0.156 | 1692<br>0.148 | 1710<br>0.148 | 1729<br>0.107 |
| 12 | 1613<br>0.152 | 1624<br>0.167 | 1641<br>0.181 | 1658<br>0.181 | 1675<br>0.163 | 1691<br>0.157 | 1709<br>0.147 | 1731<br>0.091 |
| 13 | 1614<br>0.157 | 1623<br>0.163 | 1641<br>0.180 | 1658<br>0.182 | 1675<br>0.162 | 1691<br>0.157 | 1709<br>0.144 | 1732<br>0.081 |
| 14 | 1614<br>0.171 | 1624<br>0.172 | 1641<br>0.186 | 1659<br>0.172 | 1676<br>0.154 | 1692<br>0.145 | 1710<br>0.142 | 1733<br>0.083 |
| 15 | 1614<br>0.180 | 1627<br>0.176 | 1642<br>0.182 | 1659<br>0.165 | 1676<br>0.153 | 1693<br>0.144 | 1711<br>0.144 | 1731<br>0.088 |
| 16 | 1613<br>0.188 | 1627<br>0.186 | 1642<br>0.173 | 1659<br>0.162 | 1676<br>0.148 | 1693<br>0.143 | 1711<br>0.148 | 1730<br>0.091 |
| 17 | 1613<br>0.177 | 1627<br>0.179 | 1641<br>0.176 | 1659<br>0.168 | 1676<br>0.152 | 1693<br>0.148 | 1711<br>0.148 | 1730<br>0.095 |
| 18 | 1614<br>0.171 | 1627<br>0.166 | 1641<br>0.174 | 1659<br>0.176 | 1676<br>0.161 | 1692<br>0.152 | 1710<br>0.150 | 1733<br>0.092 |
| 19 | 1616<br>0.172 | 1624<br>0.177 | 1641<br>0.174 | 1658<br>0.166 | 1676<br>0.156 | 1692<br>0.155 | 1710<br>0.151 | 1729<br>0.101 |
| 20 | 1616<br>0.213 | 1625<br>0.193 | 1641<br>0.179 | 1659<br>0.135 | 1675<br>0.132 | 1692<br>0.149 | 1710<br>0.202 | 1731<br>0.141 |

### S3: The cystine:protein intensity ratio

**S3:**
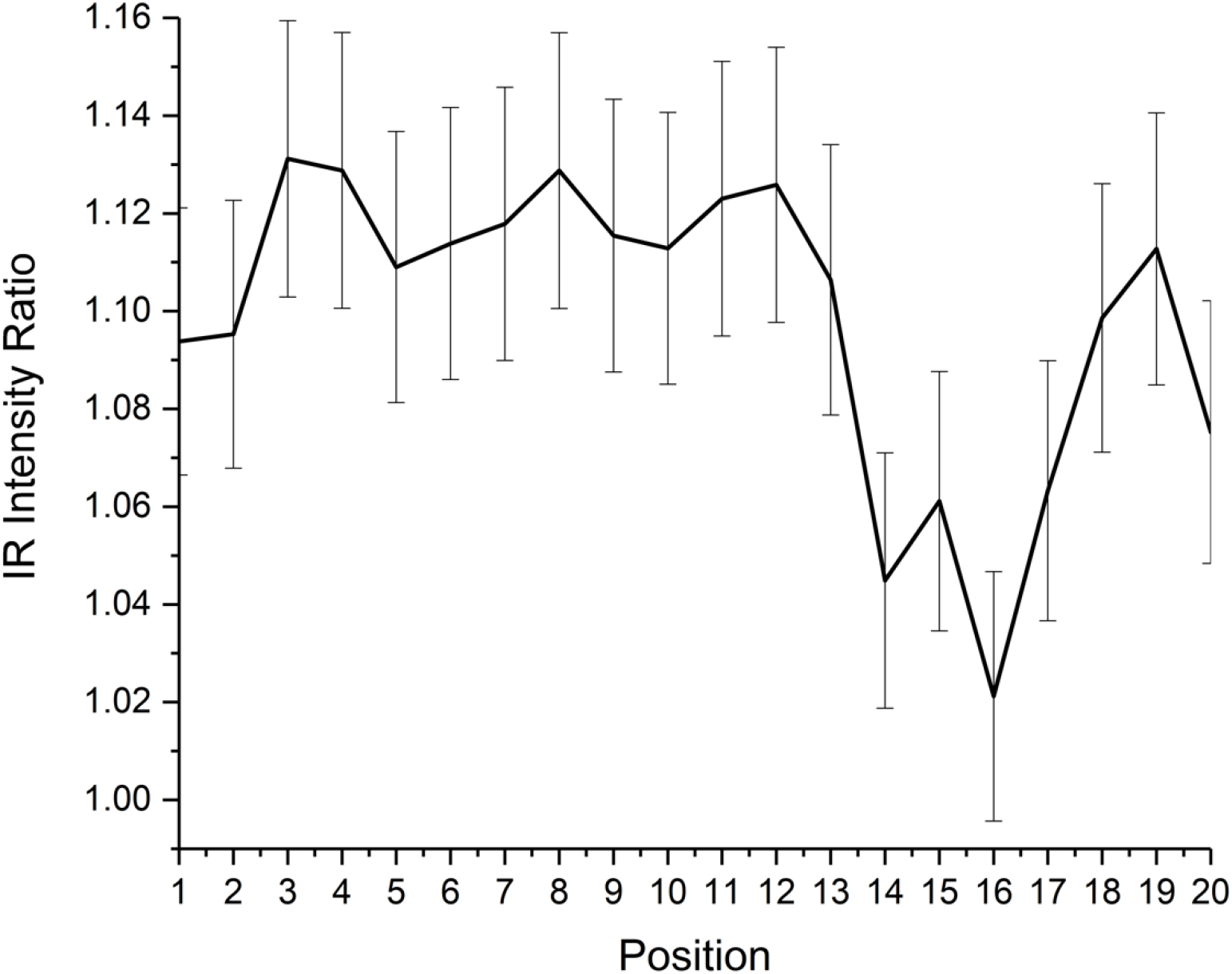
The cystine:protein intensity ratio calculated from selected sulfur-oxygen and Amide-III infrared band intensities as a function of sampling position 1 to 20.

## References

(1) Cena, K.; Clark, J. A. Thermal Insulation of Animal Coats and Human Clothing. Physics in Medicine and Biology. 1978. https://doi.org/10.1088/0031-9155/23/4/001.

(2) Randall, V. A. Hormonal Regulation of Hair Follicles Exhibits a Biological Paradox. Seminars in Cell and Developmental Biology. 2007. https://doi.org/10.1016/j.semcdb.2007.02.004.

(3) Robbins, C. R. Chemical and Physical Behavior of Human Hair, 5th Edition; 2012. https://doi.org/10.1007/9783642256110.

(4) Beigel, H. The Human Hair: Its Structure, Growth, Diseases, and Their Treatment; Henry Renshaw, 1869.

(5) Bhushan, B. Nanoscale Characterization of Human Hair and Hair Conditioners. Progress in Materials Science. 2008. https://doi.org/10.1016/j.pmatsci.2008.01.001.

(6) Wagner, R.; Joekes, I. Hair Medulla Morphology and Mechanical Properties. J Cosmet Sci 2007, 58 (4), 359–368.

(7) Rogers, G. E. Known and Unknown Features of Hair Cuticle Structure: A Brief Review. Cosmetics 2019. https://doi.org/10.3390/cosmetics6020032.

(8) Jeffrey E. Plowman and Duane P. Harland. The Hair Fibre: Proteins, Structure and Development; Springer Nature, 2018; Vol. 1054. https://doi.org/10.1007/978-981-10-8195-8.

(9) da França, S.; Dario, M.; Esteves, V.; Baby, A.; Velasco, M. Types of Hair Dye and Their Mechanisms of Action. Cosmetics 2015, 2 (2), 110–126. https://doi.org/10.3390/cosmetics2020110.

(10) Hearle, J. W. S. A Critical Review of the Structural Mechanics of Wool and Hair Fibres. Int. J. Biol. Macromol. 2000, 27 (2), 123–138. https://doi.org/10.1016/S0141-8130(00)00116-1.

(11) Wolfram, L. J. Human Hair: A Unique Physicochemical Composite. In Journal of the American Academy of Dermatology; 2003. https://doi.org/10.1067/mjd.2003.276.

(12) P. Jolles, H. Zahn, and H. H. Formation and Structure of Human Hair; Verlag, B., Ed.; 2012. https://doi.org/10.1007/978-3-0348-9223-0.

(13) Feughelman, M. Morphology and Properties of Hair. In Hair and Hair Care; 2019; pp 1–12. https://doi.org/10.1201/9780203719565-1.

(14) Swift, J. A.; Smith, J. R. Microscopical Investigations on the Epicuticle of Mammalian Keratin Fibres. J. Microsc. 2001, 204 (3), 203–211. https://doi.org/10.1046/j.1365-2818.2001.00957.x.

(15) King, N.; Bradbuby, J. The Chemical Composition of Wool V. The Epicuticle. Aust. J. Biol. Sci. 2018, 21 (2), 375. https://doi.org/10.1071/bi9680375.

(16) Bringans, S. D.; Plowman, J. E.; Dyer, J. M.; Clerens, S.; Vernon, J. A.; Bryson, W. G. Characterization of the Exocuticle A-Layer Proteins of Wool. Exp. Dermatol. 2007, 16 (11), 951–960. https://doi.org/10.1111/j.1600-0625.2007.00610.x.

(17) Rogers, G.; Koike, K. Laser Capture Microscopy in a Study of Expression of Structural Proteins in the Cuticle Cells of Human Hair. Exp. Dermatol. 2009, 18 (6), 541–547. https://doi.org/10.1111/j.1600-0625.2008.00825.x.

(18) Wei, G.; Bhushan, B.; Torgerson, P. M. Nanomechanical Characterization of Human Hair Using Nanoindentation and SEM. In Ultramicroscopy; 2005; Vol. 105, pp 248–266. https://doi.org/10.1016/j.ultramic.2005.06.033.

(19) Jones, L. N.; Rivett, D. E. The Role of 18-Methyleicosanoic Acid in the Structure and Formation of Mammalian Hair Fibres. Micron. 1997, pp 469–485. https://doi.org/10.1016/S0968-4328(97)00039-5.

(20) Dias, M. F. R. G. Hair Cosmetics: An Overview. Int. J. Trichology 2015.

(21) Wang, N.; Barfoot, R.; Butler, M.; Durkan, C. Effect of Surface Treatments on the Nanomechanical Properties of Human Hair. ACS Biomater. Sci. Eng. 2018, 4 (8), 3063–3071. https://doi.org/10.1021/acsbiomaterials.8b00687.

(22) Pudney, P. D. A.; Bonnist, E. Y. M.; Mutch, K. J.; Nicholls, R.; Rieley, H.; Stanfield, S. Confocal Raman Spectroscopy of Whole Hairs. Appl. Spectrosc. 2013. https://doi.org/10.1366/13-07086.

(23) Barton, P. M. J. A Forensic Investigation of Single Human Hair Fibres Using FTIR-ATR Spectroscopy and Chemometrics; Queensland University of Technology, Brisbane, 2011.

(24) Wei, X.; Wang, X.; Fang, Y.; Huang, Q. Comparison of Hair from Rectum Cancer Patients and from Healthy Persons by Raman Microspectroscopy and Imaging. J. Mol. Struct. 2013. https://doi.org/10.1016/j.molstruc.2013.05.005.

(25) Dazzi, A.; Prater, C. B.; Hu, Q.; Chase, D. B.; Rabolt, J. F.; Marcott, C. AFM-IR: Combining Atomic Force Microscopy and Infrared Spectroscopy for Nanoscale Chemical Characterization. Appl. Spectrosc. 2012. https://doi.org/10.1366/12-06804.

(26) Marcott, C.; Lo, M.; Kjoller, K.; Fiat, F.; Baghdadli, N.; Balooch, G.; Luengo, G. S. Localization of Human Hair Structural Lipids Using Nanoscale Infrared Spectroscopy and Imaging. Appl. Spectrosc. 2014. https://doi.org/10.1366/13-07328.

(27) Fellows, A. P.; Casford, M. T. L.; Davies, P. B. Spectral Analysis and Deconvolution of the Amide I Band of Proteins Presenting with High-Frequency Noise and Baseline Shifts. Appl. Spectrosc. 2019.

(28) Panayiotuo, H. Vibrational Spectroscopy of Keratin Fibers, A Forensic Approach, Queensland University of Technology, 2004.

(29) Qin, Z.; Buehler, M. J. Molecular Dynamics Simulation of the α-Helix to β-Sheet Transition in Coiled Protein Filaments: Evidence for a Critical Filament Length Scale. Phys. Rev. Lett. 2010, 104 (19). https://doi.org/10.1103/PhysRevLett.104.198304.

(30) Afrin, R.; Takahashi, I.; Shiga, K.; Ikai, A. Tensile Mechanics of Alanine-Based Helical Polypeptide: Force Spectroscopy versus Computer Simulations. Biophys. J. 2009, 96 (3), 1105–1114. https://doi.org/10.1016/j.bpj.2008.10.046.

(31) Song, S.-H.; Lim, J. H.; Son, S. K.; Choi, J.; Kang, N.-G.; Lee, S.-M. Prevention of Lipid Loss from Hair by Surface and Internal Modification. Sci. Rep. 2019, 9 (1), 9834. https://doi.org/10.1038/s41598-019-46370-x.

(32) Nachtigal, J.; Robbins, C. Intermediate Oxidation Products of Cystine in Oxidized Hair. Text. Res. J. 1970, 40 (5), 454–457. https://doi.org/10.1177/004051757004000511.

(33) Kogelheide, F.; Kartaschew, K.; Strack, M.; Baldus, S.; Metzler-Nolte, N.; Havenith, M.; Awakowicz, P.; Stapelmann, K.; Lackmann, J. W. FTIR Spectroscopy of Cysteine as a Ready-to-Use Method for the Investigation of Plasma-Induced Chemical Modifications of Macromolecules. J. Phys. D. Appl. Phys. 2016, 49 (8). https://doi.org/10.1088/0022-3727/49/8/084004.

(34) Dourado, A. H. B.; De Lima Batista, A. P.; Oliveira-Filho, A. G. S.; Sumodjo, P. T. A.; Cordoba De Torresi, S. I. L-Cysteine Electrooxidation in Alkaline and Acidic Media: A Combined Spectroelectrochemical and Computational Study. RSC Adv. 2017, 7 (13), 7492–7501. https://doi.org/10.1039/c6ra26576f.

(35) Singh, B. R.; DeOliveira, D. B.; Fu, F.-N.; Fuller, M. P. Fourier Transform Infrared Analysis of Amide III Bands of Proteins for the Secondary Structure Estimation. Biomol. Spectrosc. III 2005, 1890, 47–55. https://doi.org/10.1117/12.145242.

(36) Stanic, V.; Maia, F. C. B.; Freitas, R. de O.; Montoro, F. E.; Evans-Lutterodt, K. The Chemical Fingerprint of Hair Melanosomes by Infrared Nano-Spectroscopy. Nanoscale 2018, 10 (29), 14245–14253. https://doi.org/10.1039/c8nr03146k.

(37) Nyquist, R. Sulfoxides, Sulfones, Sulfates, Monothiosulfates, Sulfonyl Halides, Sulfites, Sulfonamides, Sulfonates, and N-Sulfinyl Anilines. In Interpreting Infrared, Raman, and Nuclear Magnetic Resonance Spectra; 2007; pp 85–117. https://doi.org/10.1016/b978-012523475-7/50185-6.

(38) Nikonenko, N. A.; Buslov, D. K.; Sushko, N. I.; Zhbankov, R. G. Investigation of Stretching Vibrations of Glycosidic Linkages in Disaccharides and Polysaccarides with Use of IR Spectra Deconvolution. Biopolym. – Biospectroscopy Sect. 2000. https://doi.org/10.1002/1097-0282(2000)57:4<257::AID-BIP7>3.0.CO;2-3.

(39) Higgins, H. G.; Stewart, C. M.; Harrington, K. J. Infrared Spectra of Cellulose and Related Polysaccharides. J. Polym. Sci. 1961. https://doi.org/10.1002/pol.1961.120510105.

(40) McMullen, R.; Laura, D.; Chen, S.; Koelmel, D.; Zhang, G.; Gillece, T. Determination of Physicochemical Properties of Delipidized Hair. J. Cosmet. Sci. 2013, 64, 355–370.

(41) Robbins, C. The Cell Membrane Complex: Three Related but Different Cellular Cohesion Components of Mammalian Hair Fibers. Int. J. Cosmet. Sci. 2010. https://doi.org/10.1111/j.1468-2494.2010.00577_6.x.

(42) Robinson, V. N. E. A Study of Damaged Hair. J. $oc. Cosmet. Chem 1976.

(43) Ruetsch, S. B.; Kamath, Y. K. Change in Surface Chemistry of the Cuticle of Human Hair by Chemical and Photochemical Oxidation. Int. J. Cosmet. Sci. 2005, 27 (2), 142–143. https://doi.org/10.1111/j.1467-2494.2005.00259_3.x.

(44) Okamoto, M.; Ishikawa, K.; Tanji, N.; Aoyagi, S. Investigation of the Damage on the Outermost Hair Surface Using ToF-SIMS and XPS. In Surface and Interface Analysis; 2012; Vol. 44, pp 736–739. https://doi.org/10.1002/sia.3878.

(45) Feughelman, M.; Willis, B. K. Mechanical Extension of Human Hair and the Movement of the Cuticle. J. Cosmet. Sci. 2001, 52, 185–193.

